# UNRAVELING THE LUNG VASCULAR REMODELING IN PULMONARY HYPERTENSION USING A QUANTITATIVE DIGITAL PATHOLOGY SOFTWARE

**DOI:** 10.1101/2024.07.01.601469

**Authors:** Cindy Serdjebi, Florine Chandes, Marzena Biernat, Bastien Lepoivre, Dany Salvail, Charles-E. Laurent

## Abstract

Pulmonary arterial hypertension (PAH) is a rare chronic life-threatening disorder, characterized by the elevation of the mean pulmonary arterial pressure above 20 mmHg at rest. Histologically, PAH induces lung vascular remodeling, with the thickening of vessel wall. The conventional histological analysis commonly used in non-clinical models to assess lung vascular remodeling relies on manual measurements of representative lung vessels and is time-consuming. We have developed a fully automated reader-independent software (MorphoQuant-Lung) to both specifically detect vessels and measure vascular wall components from a-SMA rat lung sections. Analysis was performed on monocrotaline-and Sugen/hypoxia-induced PH rat models, treated or not with Sildenafil. The software requires 3-5 minutes to detect up to 1500 vessels per section, classify them per size, quantify intima, media and wall thicknesses, and calculate their level of occlusion. A comparison of our digital analysis results with those of the pathologist’s conventional visual analysis was performed for wall thickness and lumen radius showing a strong correlation between the two techniques (r: 0.80 and r: 0.88) regardless of the rat model. In addition, the occlusion estimated by automated analysis also strongly correlated with the mean pulmonary arterial pressure and the pulmonary vascular resistance (r ranging from 0.71 to 0.83) in both rat models. The added value of the present digital analysis paves the way for a more in-depth understanding of the PAH physiopathology in preclinical research and provides a robust and reliable tool for efficient therapeutic drug development.

## Introduction

Pulmonary arterial hypertension (PAH) is a rare chronic life-threatening disorder characterized hemodynamically by an elevation of the mean pulmonary arterial pressure (mPAP) above 20 mm Hg at rest despite a normal pulmonary wedge pressure lower than 15 mmHg. PAH leads to right ventricular failure and eventually death (1,2). As defined by the 6^th^ World Symposium on Pulmonary Hypertension, Group 1 PAH comprises various etiologies, including idiopathic, heritable, drug-and toxin-induced PAH, but also PAH associated with other conditions (3,4). The prevalence of PAH is approximately 15 to 50 cases per million within the United States and Europe, with a marked predominance in females, while the prognosis is poorer in males (3).

Histopathology analyses reveal that PAH predominantly affects the pulmonary arterial resistance, and is characterized by an abnormal remodeling of the lung vasculature. Small and medium vessels, with a diameter up to 100 µm, are particularly affected (5–7). Among the key-pathological changes figure intimal hyperplasia, medial hypertrophy, and adventitial proliferation. Moreover, in situ thrombosis, and inflammation are observed (1). These changes lead to the progressive obliteration of vessels, eventually responsible for an increased pulmonary vascular resistance (PVR) and mPAP, right ventricle dilatation and subsequently dysfunction (1,8). Plexiform lesions, which are complex vascular formations originating from remodeled arteries, are pathognomonic features of PAH (9–11).

Several experimental animal models were developed to investigate the underlying mechanisms involved in the pulmonary hypertension (PH) pathogenesis and consequently improve therapeutic drug development (12). Among the most commonly used, the non-invasive monocrotaline (MCT)-induced PH rat model is appreciated for its ease of induction, and exhibits endothelial toxicity and marked lung inflammation (11–14). However, the pathologic features of the MCT rat model are partially reflective of the human PAH. A favored model is the Sugen5416-hypoxia (SuHx)-induced PH rat model, resulting from anti-VEGFR-2 inhibitor Sugen5416 (semaxanib) treatment combined with chronic hypoxia (15). The SuHx rat model is pathophysiologically considered as the closest preclinical model to human PAH (7,15). Upon returning to normoxic conditions following 21 days of hypoxia, damaged endothelial cells of pulmonary artery exhibit abnormal proliferation, which may conduct to the obstruction of the pulmonary artery lumen. This process can give rise to plexiform lesions.

Previous works have quantified the histological characteristics of these two PH rat models, but to a restricted extent due to the limitations of the conventional methods used. Conventional assessment of vessel morphometry consists of visually selecting a limited number of fields of view from H&E-or elastin-stained lung sections in order to manually identify blood vessels (generally up to 50) and to measure the vascular wall thickness for each vessel (2 measurements/vessel). Other also uses a-smooth muscle actin (a-SMA) labelled sections to focus on the medial hypertrophy (16,17). Additional limitations include a global analysis of endothelial cell and/or muscle cell proliferation, as well as the exclusion of blood vessels of irregular shape (non-circular). We must finally emphasize that the current conventional histological analysis is time-consuming.

Today, several medications are approved for PAH treatment, but none, including Sildenafil, reverses vascular remodeling (18). Enhancing our ability to analyze the intimate mechanisms of vascular remodeling will contribute to the development of the next generation of drugs, potentially capable of treating late-phase PAH patients at a stage where they are considered beyond reversible today. The aim of the present work was to develop a quantitative histological analysis of PH, based on a dedicated proprietary digital software (MorphoQuant-Lung), allowing fully automated, accurate and reliable measurements of morphological changes of lung vessels in rat models. Analyses were carried out on two conventional models, MCT and SuHx PH rat models treated or not with Sildenafil. Histopathological findings obtained from the present digital analysis were compared with those of the current conventional assessment and correlated with the hemodynamic parameters.

## Materials and methods

### Study Design

#### Animals

Adult male Sprague Dawley rats, weighing from 200 to 250 g at study initiation were purchased from Charles River Laboratories (St Constant, Quebec, Canada). The Institutional Animal Care and Use Committee (IACUC) of IPS Therapeutique approved the studies under ethic protocol CL20220606-03 in strict accordance with the guidelines of the Canadian Council on Animal Care (CCAC) and the Association for Assessment and Accreditation of Laboratory Animal Care (AAALAC). For both models, food and water were given ad libitum and the animals were observed on a daily basis for any changes in their behavior and general health status.

#### Monocrotaline-induced pulmonary arterial hypertension rat model

Animals received either a single subcutaneous injection of monocrotaline (AdooQ) at 60 mg/kg solution (2 mL; 30 mg/mL stock, in dimethyl sulfoxide, DMSO) or its vehicle on Day 0 before returning to their ventilated cages. The animals were then randomized among treatment groups: Sham, MCT and M-Sild (M-Sild), based on their body weight and the results of transthoracic Dopler echocardiography (at Day 7). Animals were treated by gavage with either the vehicle (water) or Sildenafil at 30 mg/kg twice a day for 21 days after randomization.

#### Sugen-hypoxia-induced pulmonary arterial hypertension rat model

On Day 0, a solution of Sugen (SU-5416, AdooQ) at 10 mg/mL in 100% DMSO was prepared. After subcutaneous administration of Sugen (20 mg/kg) in the back of the neck of the rats, the animals were housed in normobaric hypoxic chambers ventilated with an FiO_2_ equivalent to 0.10 (10%) using a mixture of nitrogen and ambient air controlled by the ventilated cage system. The animals remained under hypoxia for 21 days (3 weeks) with one bedding change per week, for limiting exposure to ambient oxygen levels for less than 10 minutes. On Day 21, animals were transferred and exposed to ambient oxygen levels until Day 56. Normoxic control animals remained in cages exposed to ambient oxygen (normoxic) levels for 56 days. The animals were randomized on Day 21 among treatment groups: normoxic (Nx), SuHx and S-Sildenafil (S-Sild), based on their body weight and the results of transthoracic Doppler echocardiography. The animals received treatment by gavage with either the vehicle (water) or Sildenafil 50 mg/kg bid from Day 22 to 55.

### Hemodynamic measurements of PH induction

On Day 28 (MCT) or Day 56 (SuHx), approximately 18-24h after the last oral dosing, the animals were anesthetized with a mixture of 2 to 2.5% isoflurane USP (Abbot Laboratories, Montreal, Canada) in oxygen, and placed on a heating pad to maintain body temperature. SpO2 were taken. The rats were tracheotomized and immediately ventilated by means of a positive-pressure rodent respirator set at ≈ 10 mL/kg body weight at a frequency of 65-70 strokes/min. A cannula connected to a pressure transducer was inserted into the left femoral artery to measure the arterial blood pressure. The heart was exposed through a sternotomy and a 20 GA Insyte was introduced into the right ventricle and rapidly hooked up to a saline filled PE-190 catheter connected to a transducer. Following a few seconds of right ventricular pressure recording, the Insyte was further advanced into the pulmonary artery to allow PAP recording for an additional 60 seconds. Hemodynamic parameters were recorded continuously for the duration of the procedure or until loss of PAP signal.

### Tissue collection and histology

Following hemodynamic monitoring, the anesthetized animals were exsanguinated and the chest cavity was further opened to expose the lungs and the heart as previously described (16). The muscle over the trachea was dissected away to harvest the lungs and heart. The collected tissues were rinsed with 15 mL of saline solution (0.9% NaCl) to remove any excess of blood. As part of the Fulton’s index, the heart was dissected to separate the right ventricle from the left ventricle with septum, and then weighed separately. The left lung lobe was then inflated using a 60 mL syringe filled with fixative (10% neutral buffered formalin at a 1:20 tissue to fixative ratio). The tissues were kept in formalin for 24-48 hours before being transferred in 70% ethanol and then embedded in paraffin blocks.

For each animal, two replicates of three transversal consecutive sections (5-µm thick) were cut, each replicate spaced by 50 µm (Wax-it Histology services Inc., Vancouver BC. Canada). The two replicates were laid on a same slide and stained with either hematoxylin-eosin (H&E) for the conventional visual assessment, Van-Gieson as a specific elastin stain or labelled with alpha-smooth muscle actin (a-SMA, Cat#M0851, Dako, 1:1000) as a specific marker for muscle. Whole slides were thus scanned at the magnification of either 20 or 40X on a Leica Aperio AT2 scanner and converted as whole slide images into .svs format image files. Images with each histology techniques were tested for the development of an automated tool to assess the lung vascular remodeling.

### Conventional assessment of vascular remodeling in PH

For the conventional evaluation of the lung vascular remodeling, H&E-stained sections were scanned at 40X and used as described previously (17). Briefly, the pulmonary arteries/arterioles visualized from about 20 random fields of view were identified and categorized according to their Feret diameter into small (10–50 µm), medium (51–100 µm), and large diameter vessels (>100 µm); and the percentage of these pulmonary blood vessels demonstrating the presence of a muscular (completely surrounded by a smooth muscle layer, > 90% circumference), semi-muscular (incompletely surrounded by a smooth muscle layer, 10 – 90% circumference) or a non-muscular (no apparent smooth muscle layer, < 10% circumference) appearance were quantified. The lumen diameter was measured from inner edge of the lamina elastic interna on one side to the other side. The lamina elastic interna is defined as the layer just below the endothelial cells. From these measurements, total vessel diameter, medial wall thickness and percentage occlusion may be determined. All analyses were carried out in a blinded fashion.

### Development of an automated software for the assessment of the lung vascular remodeling in PH

Histologically, PAH is characterized by the thickening of the medial layer mainly in pulmonary arteries and remodeling of vessel endothelial cells. Consecutive lung sections from animals from Sham, MCT, Normoxic, and SuHx groups obtained from previous in vivo experiments were stained with H&E, Van Gieson, and a-SMA IHC to investigate the most suitable histology technique to identify and quantify vessels reliably. Our initial objectives were to focus on arteries and to quantify the overall wall thickness, but also to set apart the intima from the media, given that histological responses to MCT and SuHx induction may vary and to better understand the effect of Sildenafil. H&E staining, which does not allow to reliably discriminate the intima from the media was rapidly deemed non suitable for further automated analysis. Van Gieson stain, which permits to visualize the elastin and thus the two different layers, was tried. However, our internal tests did not show significant change in colorimetry in comparison with the alveolar wall to allow reliable detection of vessels. In the context of lung tissue, a-SMA IHC specifically labels smooth muscle cells from bronchi and vessels. However, it does not allow to reliably identify arteries from veins, as some veins could exhibit muscle and some small arteries are non-muscular. We thus assumed that the changes seen in the a-SMA labelled vessels, i.e. the muscular and semi-muscular vessels, regardless of their nature (artery, vein or lymphatic), were reflective of the overall changes in arteries in the context of PH.

Therefore, we used whole slide images of a-SMA labelled sections of rat left lobe lung cut transversely. After the automated delineation of entire lung sections, size of sections (in mm^²^) and a-SMA proportionate area (expressed in percent) were measured (Fig. 1A). To identify vessels, the stained tissue was first segmented and extracted to obtain a mask. Within this tissue mask, all the delineated airspaces (several thousands) were labelled as vessels or others, and a set of descriptors was computed for each of them. We then trained a C5 algorithm model to generate a set of weighted rules allowing the automated recognition of vessels in the newly submitted whole slides images of a-SMA labelled rat lung sections.

**Figure 1.**
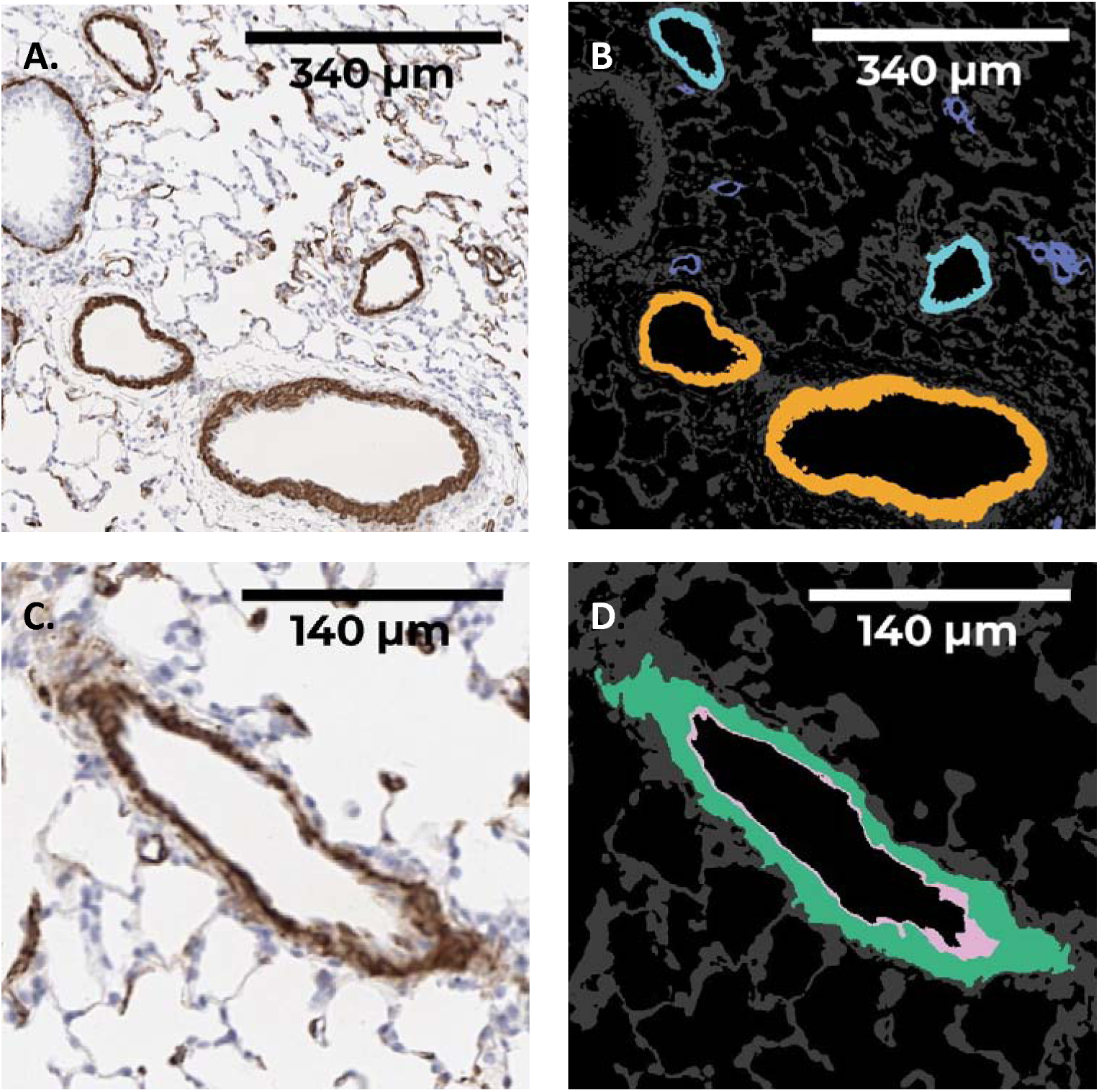
Original scans of a-SMA labelled rat lung sections and the related computational illustrations from the automated digital analysis. **a)** Original a-SMA labelled rat lung section. **b)** Mapped image of the automatically detected vessels per size category (dark blue, light blue and orange represent small, medium and large vessels, respectively). **c)** Original image of a longitudinal vessel with its a-SMA positive media layer. **d)** Processed image showing the detected vessel and its related intima (pink) and media (green) layers.

After computation of the selection of descriptors and application of the defined set of weighted rules, the actual vessels were identified and categorized according to their minimal Feret diameter (FD) as small, medium and large vessels (20 < FD ≤ 50; 50 < FD ≤ 100; 100 < FD ≤ 200 µm), respectively (Fig. 1B and 1C). As the vessels with diameter larger than 200 µm are less affected by PH vascular remodeling (2,5–7), they were not considered in the analysis. Then, for each vessel, the lumen was defined as the empty area within the delineated airspace, the intima as the non a-SMA stained tissue, and the media as the a-SMA positive layer (Fig. 1D). The area of each component (lumen, intima and media) was measured and the vessels were assumed perfectly circular to translate areas into thicknesses. Up to 1,500 muscularized vessels were detected per lung section, and 11 readouts (Table 1) were estimated for each of them, namely: vessel area, minimal Feret diameter, vessel radius, luminal area and radius, intimal area and thickness, media area and thickness, wall area and thickness (sum of intima and media), occlusion rate. Vessels were considered obstructed when the ratio wall area/vessel area was superior to 0.5. The proportion of obstructed vessels per section was also calculated. The run time analysis ranged from 3 to 5 min per section. The results presented further down were 100 % repeatable after re-analysis of all the whole slide images (data not shown).

**Table 1.** Readouts provided by automated digital analysis from a-SMA labelled rat lung sections. The readouts are provided either at the whole section level, or for each individual vessel, allowing to calculate the mean readouts per size category.

| Readouts | Per vessel | Per size category | Whole section |
| --- | --- | --- | --- |
| Section area (mm<sup>2</sup>)<br>Area covered by the lung section after automated delineation. |  |  | x |
| a-SMA proportionate area (%)<br>Area covered by a-SMA labelling and normalized versus the entire lung section area. |  |  | x |
| Total vessels (n/mm<sup>2</sup>)<br>Total number of detected muscularized blood vessels, with a small Feret diameter inferior to 200 µm, across the whole section. |  | x | x |
| Hypercellular lesions (n/mm<sup>2</sup>) |  |  | x |
| Vessel area (µm<sup>2</sup>)<br>Total area of the detected blood vessel, including lumen, intima, and media. | x |  |  |
| Minimal Feret diameter<br>Measure of the smallest diameter of a blood vessel. | x |  |  |
| Vessel radius (µm)<br>Area-extrapolated radius of the considered blood vessel after assuming it was perfectly circular. | x |  |  |
| Luminal area (µm<sup>2</sup>)<br>Area covered by the considered lumen. | x |  |  |
| Luminal radius (µm)<br>Area-extrapolated radius of the lumen considering perfectly circular blood vessels. | x | x |  |
| Intimal area (µm<sup>2</sup>)<br>Area covered by the intima layer within a considered blood vessel. | x |  |  |
| Intimal thickness (µm)<br>Area-extrapolated thickness of the intima layer | x | x |  |
| Medial area (µm<sup>2</sup>)<br>Area covered by the media layer within a considered blood vessel. | x |  |  |
| Medial thickness (µm)<br>Area-extrapolated thickness of the media layer, assuming perfectly circular blood vessels. | x | x |  |
| Wall thickness (µm)<br>Thickness of the wall (intima + media) | x | x |  |
| Occlusion (%)<br>The occlusion is the ratio of the vessel wall over the total radius of the vessel (lumen radius + wall) | x | x |  |
| Occluded vessel proportion (%)<br>A vessel is considered occluded if its obliteration is equal or |  | x | x |
| superior to 50%. Occluded vessel proportion is the ratio of the number of occluded vessels versus the total number of detected muscularized vessels. |  |  |  |

### Statistical analysis

For each lung section, an arithmetic mean of the computed readouts was calculated for each vessel size category and used for comparison among groups to assess correct induction and sildenafil effect on vascular remodeling. Sham and M-Sild was compared to MCT and Normoxic and S-Sildenafil was compared to SuHx. After checking the normality distribution of each group, using the appropriate t-test (t-test or Mann-Whitney) according to the normality distribution assumption. Differences were considered significant when the p-value was lower than 0.05. Correlations between the visual assessment and quantitative data from MorphoQuant were also investigated by calculating the Pearson’s correlation for wall thickness, lumen radius and obstruction, gathering data from all size categories. Correlations with the mPAP and PVR were calculated considering one overall wall thickness measurement, considering the proportion of vessels for each category, and one measurement of mPAP and PVR per subject.

## RESULTS

### Automated digital analysis of vascular remodeling in MCT-induced PH rats

Following a single subcutaneous injection of MCT at 60 mg/kg on Day 0, the animals underwent pulmonary hemodynamic measurements and cardiac function assessments on Day 28. Although there were no significant physiological changes in the mean systemic arterial pressure (mSAP) observed among Sham, MCT, and M-Sild groups, the mean pulmonary arterial pressure (mPAP) and Fulton’s index demonstrated a significant 2.2-fold and 2.1-fold increase, respectively, in the MCT group, indicating the successful induction of PH (Fig. 2A, 2B, and 2C). The rise in mPAP, accompanied by remodeling of the pulmonary vasculature, results in an increase in PVR (Fig. S1). This, in turn, elevates cardiac afterload, ultimately culminating in right ventricular heart failure, as indicated by the diminished cardiac function parameters observed in the MCT group. These parameters include reductions in stroke volume, cardiac output, and Vmax (Fig. S1 and Fig. 2D). Upon treatment with Sildenafil at a dose of 30 mg/kg administered twice daily via oral gavage, starting from Day 7, the mPAP exhibited significant reductions of 25% compared to the untreated MCT group. This suggests a vasodilatory effect of Sildenafil with a recovery of 46.9 % (Fig. 2B). The three-week treatment with Sildenafil also contributed to a trending improvement in cardiac remodeling and functions (Fig. 2C, D and Fig. S1).

**Figure 2.**
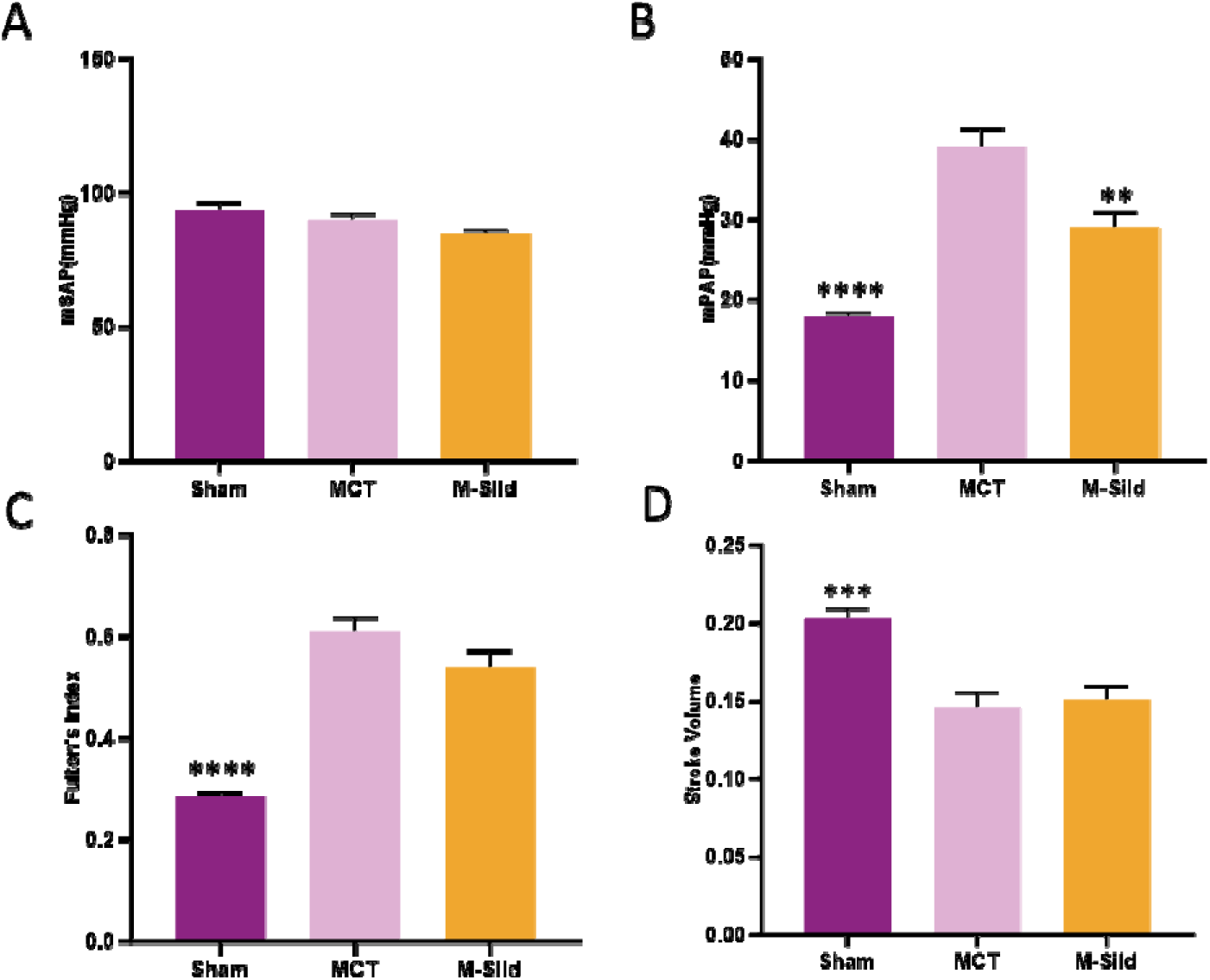
Hemodynamic measurements and cardiac function assessment in MCT-induced PAH rats. **a)** Mean systemic arte ial pressure. **b)** Mean pulmonary arterial pressure. **c)** Fulton’s index. **d)** Pulmonary valve peak velocity. The statistical analysis was performed with either a t-t st or a Mann-Whitney to compare Sham or M-Sild to MCT. MCT: monocrotaline; Sild: Sildenafil. *: p < 0.05; **: p < 0.01; ***: p < 0.001; ****: p < 0.0001.

Vascular remodeling was measured through the conventional visual assessment from H&E digitized sections on one hand and the developed automated software from a-SMA labelled sections on the other hand (Table 2). According to the histological pathologist’s evaluation based on H&E sections, there was a notable increase in wall thickness of small (2.4-fold), medium (2-fold), and large-sized vessels (1.3-fold) in pulmonary arteries from MCT animals. Animals treated with Sildenafil showed a slight, but significant reversal or prevention of the wall thickness in small (1.4-fold) and medium vessels only (1.1-fold). Consequently, the percentage of vessels categorized as muscular also significantly increased (from 9.50 to 78.33 %) while the non-muscular proportion decreased (from 81.82 to 14.57%). Animals treated with Sildenafil also demonstrated a significant reduction (1.5-fold) in muscular vessels (Fig. S2).

**Table 2.**
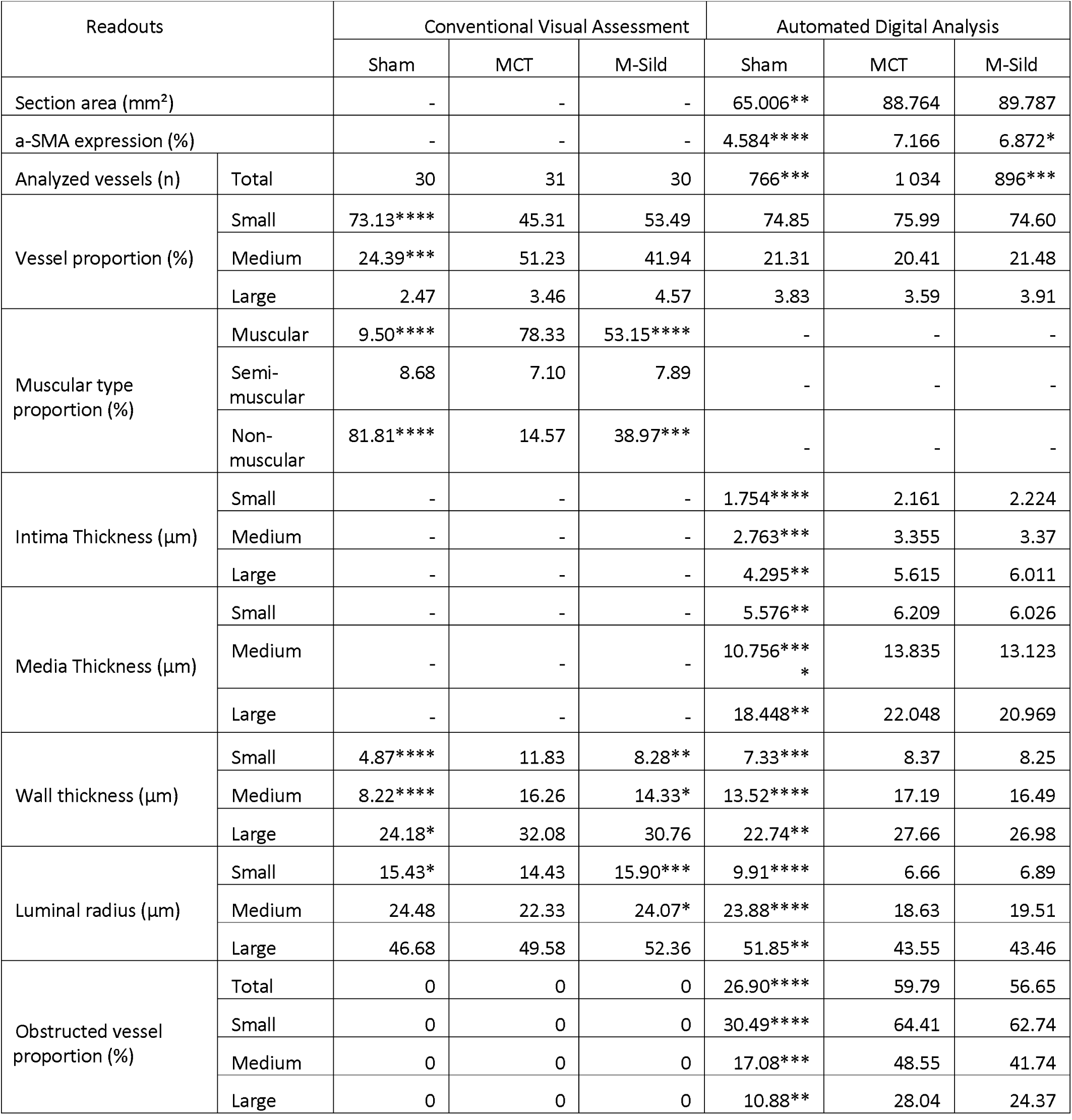
Histological characteristics of vessels in MCT rat model measured by conventional visual and automated digital analysis. Parameters labelled with “-“ were not provided by the considered analysis of PH slides. The statistical analysis was performed with either a t-test or a Mann-Whitney to compare Sham or M-Sild to MCT. MCT: monocrotaline; Sild: Sildenafil. *: p < 0.05; **: p < 0.01; ***: p < 0.001; ****: p < 0.0001.

| Readouts |  | Conventional Visual Assessment |  |  | Automated Digital Analysis |  |  |
| --- | --- | --- | --- | --- | --- | --- | --- |
|  |  | Sham | MCT | M-Sild | Sham | MCT | M-Sild |
| Section area (mm<sup>2</sup>) |  | - | - | - | 65.006** | 88.764 | 89.787 |
| a-SMA expression (%) |  | - | - | - | 4.584**** | 7.166 | 6.872* |
| Analyzed vessels (n) | Total | 30 | 31 | 30 | 766*** | 1 034 | 896*** |
| Vessel proportion (%) | Small | 73.13**** | 45.31 | 53.49 | 74.85 | 75.99 | 74.60 |
|  | Medium | 24.39*** | 51.23 | 41.94 | 21.31 | 20.41 | 21.48 |
|  | Large | 2.47 | 3.46 | 4.57 | 3.83 | 3.59 | 3.91 |
| Muscular type proportion (%) | Muscular | 9.50**** | 78.33 | 53.15**** | - | - | - |
|  | Semi-muscular | 8.68 | 7.10 | 7.89 | - | - | - |
|  | Non-muscular | 81.81**** | 14.57 | 38.97*** | - | - | - |
| Intima Thickness (µm) | Small | - | - | - | 1.754**** | 2.161 | 2.224 |
|  | Medium | - | - | - | 2.763*** | 3.355 | 3.37 |
|  | Large | - | - | - | 4.295** | 5.615 | 6.011 |
| Media Thickness (µm) | Small | - | - | - | 5.576** | 6.209 | 6.026 |
|  | Medium | - | - | - | 10.756***<br>* | 13.835 | 13.123 |
|  | Large | - | - | - | 18.448** | 22.048 | 20.969 |
| Wall thickness (µm) | Small | 4.87**** | 11.83 | 8.28** | 7.33*** | 8.37 | 8.25 |
|  | Medium | 8.22**** | 16.26 | 14.33* | 13.52**** | 17.19 | 16.49 |
|  | Large | 24.18* | 32.08 | 30.76 | 22.74** | 27.66 | 26.98 |
| Luminal radius (µm) | Small | 15.43* | 14.43 | 15.90*** | 9.91**** | 6.66 | 6.89 |
|  | Medium | 24.48 | 22.33 | 24.07* | 23.88**** | 18.63 | 19.51 |
|  | Large | 46.68 | 49.58 | 52.36 | 51.85** | 43.55 | 43.46 |
| Obstructed vessel proportion (%) | Total | 0 | 0 | 0 | 26.90**** | 59.79 | 56.65 |
|  | Small | 0 | 0 | 0 | 30.49**** | 64.41 | 62.74 |
|  | Medium | 0 | 0 | 0 | 17.08*** | 48.55 | 41.74 |
|  | Large | 0 | 0 | 0 | 10.88** | 28.04 | 24.37 |

The automated analysis showed major morphological changes between Sham and MCT animals, in relation with PH progression and vascular remodeling (Table 2). Lung sections were significantly larger (1.4-fold) with a higher a-SMA expression (1.6-fold), and a tendency to an increase of the number of muscular vessels, especially for small vessels. The luminal radius of vessels was significantly smaller for all size categories, with concomitantly a significant increase of intima and media thickness. This vascular remodeling led to a significant increase in the number of small-, medium-and large-occluded vessels (occlusion superior or equal to 50%). With respect to subjects treated with Sildenafil, a 10.9% reduction in the number of vessels was shown (p = 0.0926) compared to the MCT group, with no further change seen, suggesting that sildenafil has a limited effect on lung vasculature remodeling in MCT-induced PH rats.

### Automated digital analysis of vascular remodeling in SuHx-induced PH rats

Animals receiving 20 mg/kg of Sugen5416 were exposed to three weeks of hypoxia followed by five weeks of normoxia, were then anesthetized and instrumented to assess hemodynamic changes occurring within this model. As anticipated, the mSAP showed no significant change (Fig. 3A). In contrast, the mPAP experienced a substantial 2.9-fold increase in the SuHx group compared to the control normoxic group (Fig. 3B). Likewise, the Fulton’s Index, which measures ventricular hypertrophy, exhibited a 2.2-fold increase in the SuHx group (Fig. 3C). This shift indicated the presence of ventricular hypertrophy attributable to prolonged ventricular afterload triggered by the elevated mPAP and the 5.5-fold elevated PVR (as shown in Fig. S3). Furthermore, a comparison between the normoxic and SuHx groups demonstrated a significant decline in cardiac functions as shown by reduced stroke volume, Vmax and cardiac output (Fig. 3D, and Fig. S2). After 5-week treatment with Sildenafil at 50 mg/kg bid by oral gavage from Day 21, the mPAP showed a significant reduction, with a recovery of 50%. Cardiac function parameters also improved significantly (Fig. 3D and Fig. S3).

**Figure 3.**
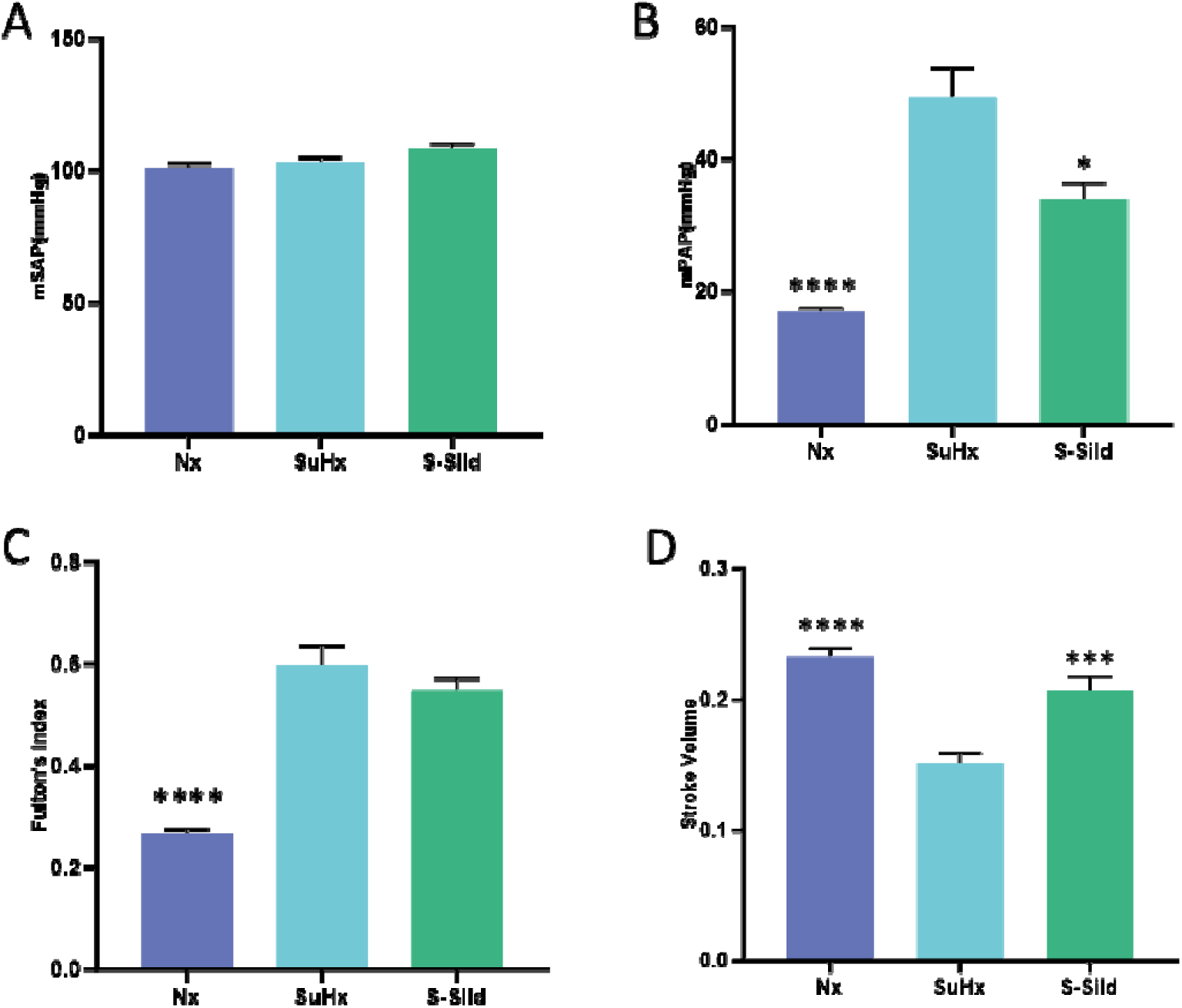
Hemodynamic measurements and cardiac function assessment in SuHx-induced PAH rat model. **a)** Mean systemic arterial pressure. **b)** Mean pulmonary arterial pressure. **c)** Fulton’s index. **d)** Pulmonary valve peak velocity. The statistical analysis was performed with either a t-test or a Mann-Whitney to compare Nx or S-Sild to SuHx. SuHx: Sugen-hypoxia; Sild: Sildenafil. *: p < 0.05; **: p < 0.01; ***: p < .001; ****: p < 0.0001.

When vascular remodeling was reported using the visual assessment of H&E sections from SuHx animals, the pathologist’s evaluation showed a significant increase in wall thickness of small (2.2-fold), medium (1.9-fold) and large (1.1-fold) pulmonary arteries (Table 3). Consequently, the proportion of lung arteries categorized as muscular significantly increased (from 15.23 to 84.35 %) while the proportion of non-muscular vessels decreased (from 71.30 to 10.63 %). Animals treated with Sildenafil showed a slightly, but significant reduction of wall thickness in small (1.4-fold) and medium (1.2-fold) vessels with concomitantly a significant decrease (1.4-fold) of the percentage of muscular vessels (Fig. S4).

**Table 3.**
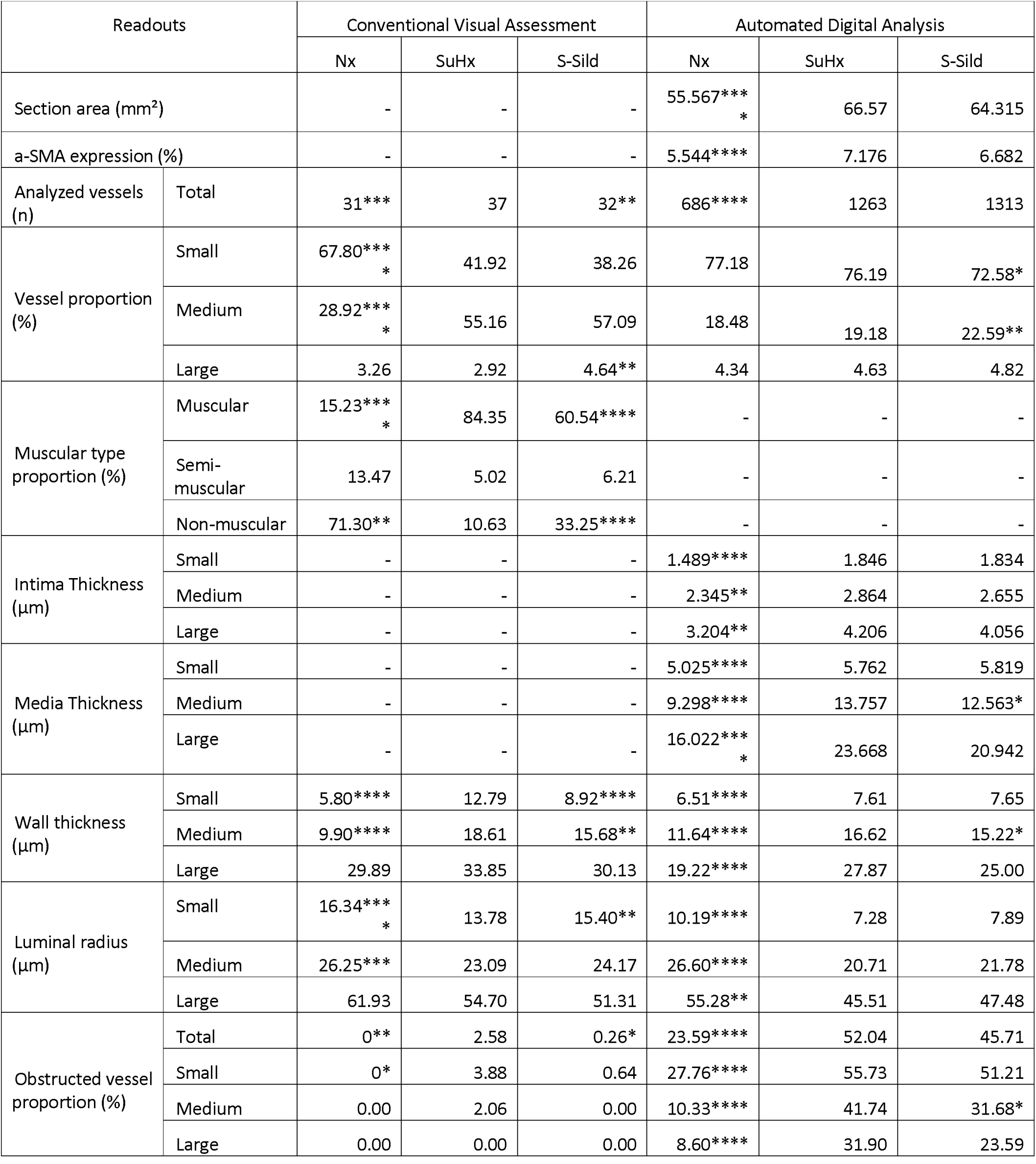
Histological characteristics of vessels in MCT rat model measured by conventional visual and automated digital analysis. Parameters labelled with “-“ were not provided by the considered analysis of PH slides. The statistical analysis was performed after checking normality assumptions. Either a t-test or a Mann-Whitney was performed to compare Nx or S-Sild to SuHx + Veh. Nx: Normoxic; SuHx: Sugen-hypoxia; Veh: vehicle; Sild: Sildenafil. *: p < 0.05; **: p < 0.01; ***: p < 0.001; ****: p < 0.0001.

| Readouts |  | Conventional Visual Assessment |  |  | Automated Digital Analysis |  |  |
| --- | --- | --- | --- | --- | --- | --- | --- |
|  |  | Nx | SuHx | S-Sild | Nx | SuHx | S-Sild |
| Section area (mm<sup>2</sup>) |  | - | - | - | 55.567***<br>* | 66.57 | 64.315 |
| a-SMA expression (%) |  | - | - | - | 5.544**** | 7.176 | 6.682 |
| Analyzed vessels (n) | Total | 31*** | 37 | 32** | 686**** | 1263 | 1313 |
| Vessel proportion (%) | Small | 67.80***<br>* | 41.92 | 38.26 | 77.18 | 76.19 | 72.58* |
|  | Medium | 28.92***<br>* | 55.16 | 57.09 | 18.48 | 19.18 | 22.59** |
|  | Large | 3.26 | 2.92 | 4.64** | 4.34 | 4.63 | 4.82 |
| Muscular type proportion (%) | Muscular | 15.23***<br>* | 84.35 | 60.54**** | - | - | - |
|  | Semi-muscular | 13.47 | 5.02 | 6.21 | - | - | - |
|  | Non-muscular | 71.30** | 10.63 | 33.25**** | - | - | - |
| Intima Thickness (μm) | Small | - | - | - | 1.489**** | 1.846 | 1.834 |
|  | Medium | - | - | - | 2.345** | 2.864 | 2.655 |
|  | Large | - | - | - | 3.204** | 4.206 | 4.056 |
| Media Thickness (μm) | Small | - | - | - | 5.025**** | 5.762 | 5.819 |
|  | Medium | - | - | - | 9.298**** | 13.757 | 12.563* |
|  | Large | - | - | - | 16.022***<br>* | 23.668 | 20.942 |
| Wall thickness (μm) | Small | 5.80**** | 12.79 | 8.92**** | 6.51**** | 7.61 | 7.65 |
|  | Medium | 9.90**** | 18.61 | 15.68** | 11.64**** | 16.62 | 15.22* |
|  | Large | 29.89 | 33.85 | 30.13 | 19.22**** | 27.87 | 25.00 |
| Luminal radius (μm) | Small | 16.34***<br>* | 13.78 | 15.40** | 10.19**** | 7.28 | 7.89 |
|  | Medium | 26.25*** | 23.09 | 24.17 | 26.60**** | 20.71 | 21.78 |
|  | Large | 61.93 | 54.70 | 51.31 | 55.28** | 45.51 | 47.48 |
| Obstructed vessel proportion (%) | Total | 0** | 2.58 | 0.26* | 23.59**** | 52.04 | 45.71 |
|  | Small | 0* | 3.88 | 0.64 | 27.76**** | 55.73 | 51.21 |
|  | Medium | 0.00 | 2.06 | 0.00 | 10.33**** | 41.74 | 31.68* |
|  | Large | 0.00 | 0.00 | 0.00 | 8.60**** | 31.90 | 23.59 |

Vascular remodeling changes were also assessed in SuHx animals by automated digital analysis (Table 3). In this model, lung section area and a-SMA expression were significantly increased (1.3-fold) in comparison with normoxic controls. There were more detected muscularized vessels in PH-induced animals (1.8-fold), impacting all size categories. The intima and media thickness increased significantly whatever the vessel size, with consequently a significant increase of the wall thickness in small (1.2-fold), medium (1.4-fold) and large (1.5-fold) vessels. Accordingly, the luminal radius decreased, leading to a significant increase of small-(2-fold), medium-(4-fold) and large-(3.7-fold) occluded vessels. In SuHx animals, Sildenafil induced a significant reduction of media in medium vessels (1.3-fold), and a tendency was also observed in large vessels.

### Comparisons and correlations of between visual assessment, digital quantification and hemodynamic testing

The analysis times were compared between the visual assessment and the digital analysis processes: visual assessment took between 30 to 40 minutes per lung section to manually select and measure pulmonary arteries versus 3 to 5 minutes with the automated digital analysis hence reducing 6-to 13-fold the analysis time (data not shown). In addition, the number of vessels considered during the automated digital analysis was higher (up to 35-fold) than that of the visual assessment, regardless of the size category and animal models (Table 2 and 3).

The digital quantification showed significant increase of the number of detected vessels in MCT and SuHx groups in comparison to control animals (Fig. 4). With respect to the vessel wall, the automated digital program was able to separate and measure both intima and media layers. In contrast, an overall assessment of the wall thickness was provided by the pathologist. Regardless of vessel size and animal model, the digital quantification and the visual assessment produced a good correlation of 0.79 (p-value < 0.0001, Fig. 5). A similar finding was observed for the luminal radius, where the correlation between manual visual and digital assessment reached 0.72 (p-value < 0.0001, Fig. S5) while the correlation was moderate for vessel occlusion (r = 0.49, p < 0.0001, Fig. S6). Of note, the digital analysis does not classify vessels into non-, semi-and muscular vessels.

**Figure 4.**
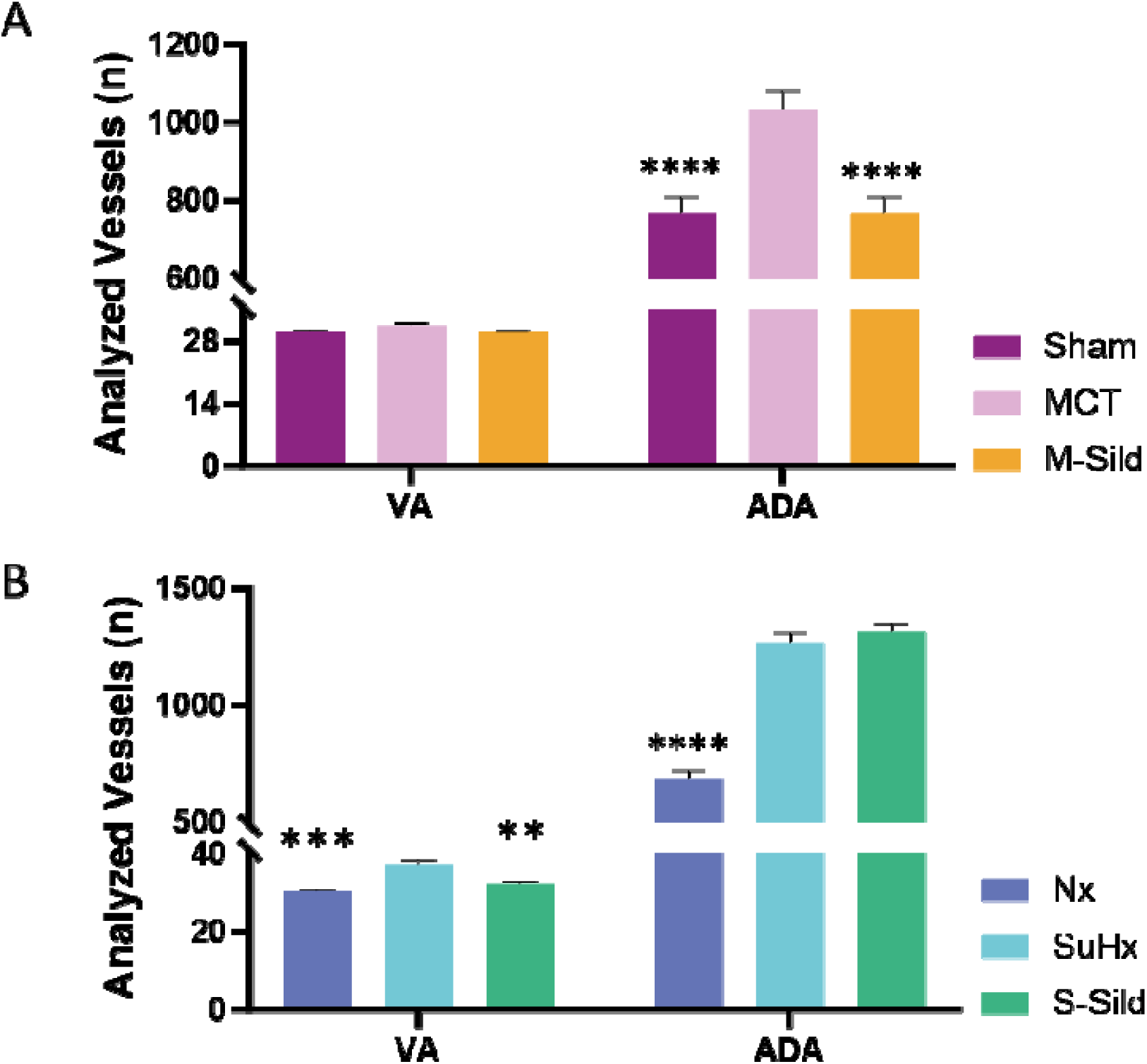
Number of analyzed vessels per rat lung section b assessment and automated digital analysis. **a)** Number of vessels in the MCT experiment. **b)** Number of analyzed vesse SuHx experiment. The statistical analysis was performed with t-test or a Mann-Whitney to compare Nx or S-Sild to SuHx hand and Sham or M-Sild to MCT on the other hand. V assessment; ADA: automated digital analysis. Nx: Normoxi Sugen-hypoxia; MCT: Monocrotaline; Sild: Sildenafil. *: p < 0.< 0.01; ***: p < 0.001; ****: p < 0.0001.

**Figure 5.**
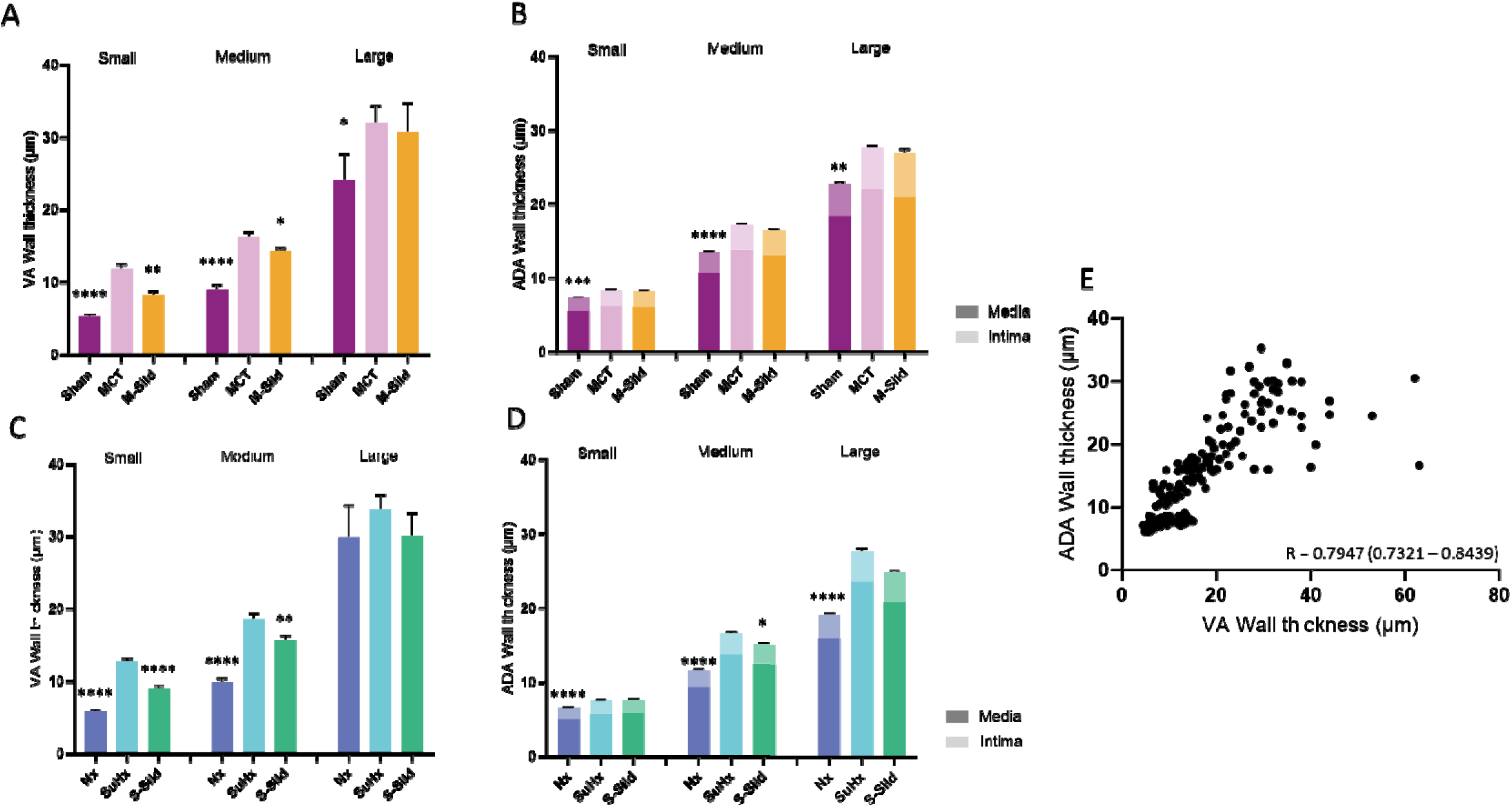
Comparison of wall thickness between visual assessment and automated digital quantification in MCT-and SuHx-induced PAH all thickness measured by VA **(a)** and ADA **(b)** in the MCT experiment. Wall thickness assessed by VA **(c)** and ADA **(d)** in the SuHx ment. In panels B and D, the bottom of the data columns (dark) illustrates media thicknesses, while the top of the column (lighter illustrates the intimal thickness. VA: visual assessment; ADA: auto ated digital analysis; Nx: Normoxic; SuHx: Sugen-hypoxia; MCT: rotaline; Sild: Sildenafil. *: p < 0.05; **: p < 0.01; ***: p < 0.001; ****: p < 0.0001. Comparison with either Vehicle groups.

Lastly, we assessed whether the vascular remodeling quantified by automated morphometric analysis aligned with the hemodynamic findings, especially the mPAP and PVR; mPAP is the principal clinical criterion for PH and PVR drives the prognosis. So, we associated mPAP and PVR with mean obstruction rates and obtained good correlations with r = 0.82 and 0.75 for mPAP and PVR, respectively in MCT animals and r = 0.80 and 0.71 for mPAP and PVR, respectively in SuHx (p-value < 0.0001 for all, Fig. 6). This supports the hypothesis that vascular remodeling-related occlusion leads to the increase of hemodynamic pressure.

**Figure 6.**
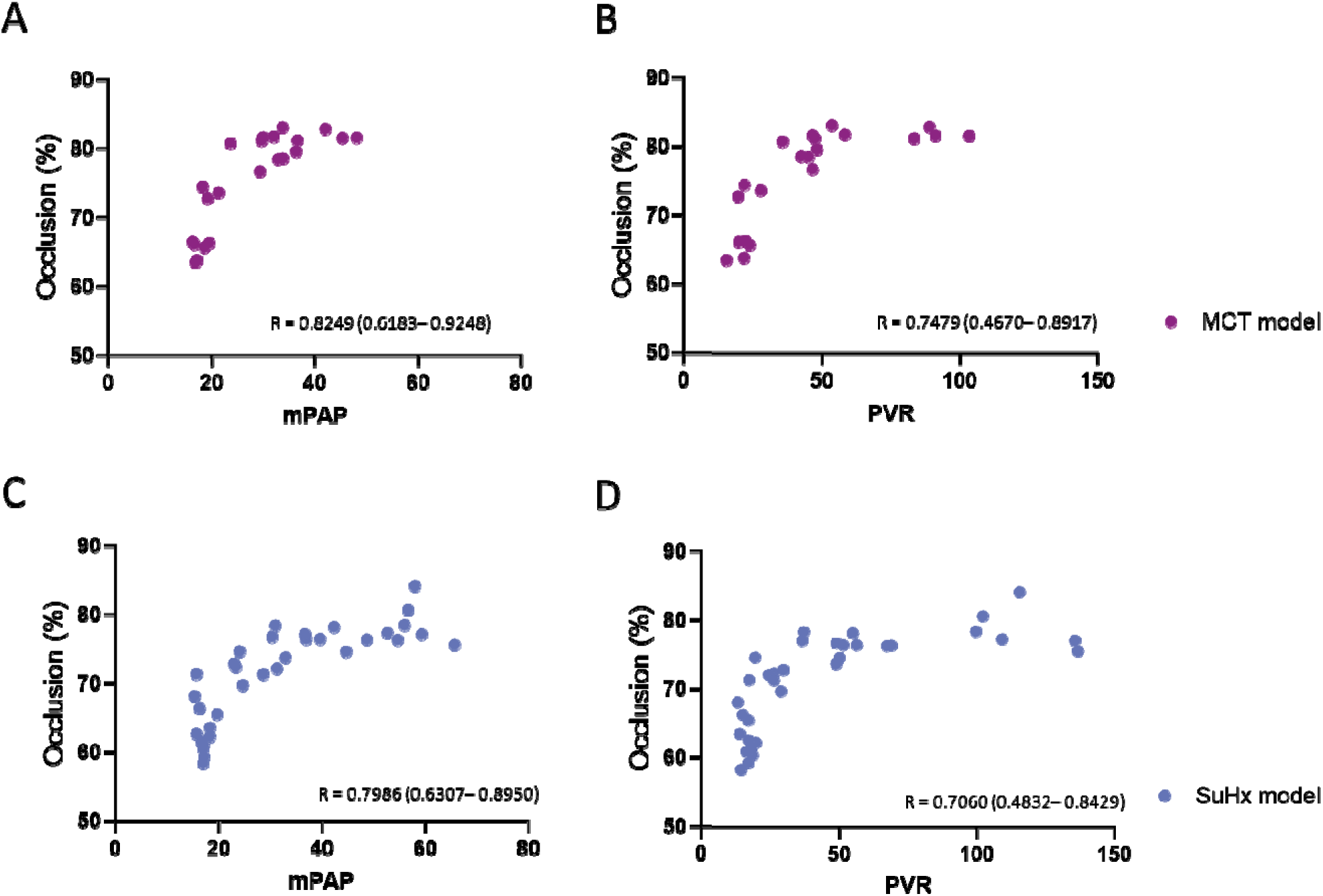
Correlation of mean occlusion measured by automated digital analysis with mPAP and PVR in MCT-and SuHx-ind ced PAH rats. mPAP: mean pulmonary arterial pressure; PVR: pulmonary vascular resistance.

## DISCUSSION

We developed a fully automated user-independent software for the comprehensive analysis of lung vascular remodeling in the two most commonly used PH rat models, i.e. MCT and SuHx rat models.

For this purpose, a-SMA immunolabeled lung sections were chosen instead of the conventional H&E or elastin stain in order to facilitate lung vessel detection and visualization of the tunica intima and tunica media (16, 19, 20). The present software allowed a significant reduction of the time necessary to process a lung section, improving it by 6-to 13-fold.

Despite the present analysis focusing on a-SMA-delineated vessels, which correspond to muscular or semi-muscular vessels, the software allowed a comprehensive detection of vessels (up to 1,500 vessels), regardless of their size or nature. A total of 37 readouts were processed and calculated over each whole section. Moreover, the vessels were automatically categorized into small, medium and large vessels according to their diameter, further defining the extent of vascular remodeling. Given that PH and PAH are generally considered pathologies of the small pulmonary vasculature, segregating the data per vessel size consequently provides what is expected to be greater predictivity of therapeutic drug efficacy for preclinical candidates. In our opinion, one of the most significant advances from this work is to measure intima and media thickness separately from the same lung section, to provide a unique resolution into potential mechanisms of efficacy – such as inhibition of endothelial or vascular smooth muscle proliferation. Usually, Von-Willebrand factor or CD31 immunolabeled slides are necessary to measure specifically intima (21), requiring the use of consecutive slides or dual staining for the assessment of intima and media altogether, which may hamper accuracy. As vessel occlusion generally results from either intima and/or media thickening in MCT-and SuHx-induced PH models (7, 12), providing this level of resolution contributes to a deeper understanding of the mechanisms underlying vascular remodeling and a major help for the development of new non-clinical PH models (22, 23). However, since this version of the software uses a-SMA labeled section, it does not “see” non-muscular vessels. Furthermore, neither information is given on the nature of the vessel (vein or artery, for instance) nor is the level of “muscularization” categorized in the conventional manner (non-muscular, semi-muscular, or fully muscular).

The present digital analysis captures significant changes in lung vasculature of two PH rat models, across nearly all the developed readouts: increase in the number of detected vessels, thickening of the intima, media and wall, smaller luminal radius, and greater proportion of obstructed vessels. Digital analysis did not detect changes in the size distribution of vessels, contrary to conventional visual assessment. This raises interesting questions regarding the roles and becoming of non-muscular vessels during PAH development or angiogenesis which would imply an increase of the proportion of small vessels (24–27).

The effects of Sildenafil, used as a positive control, were investigated in both models by manual visual assessment as well as by automated digital analysis. In the MCT experiment, automated analysis demonstrated that Sildenafil treatment resulted in a significant decrease of the number of lung vessels, with no further remodeling. In SuHx, while no change was observed in the number of vessels, Sildenafil-treated animals exhibited a slightly but significant thinner media and wall for medium vessels, as well as a decrease of obstructed medium-sized vessels. This suggests that the vasodilative effects of Sildenafil, although of significant impact on hemodynamic and cardiovascular functions, remain modest on the prevention or reduction of the lung vascular remodeling in the two rat models studied (28). This is also coherent with the fact that Sildenafil is administered relatively early in the MCT experiment (on Day 7), when the disease just provides mild signs of cardiac dysfunction, suggesting that it may slow down the disease progression and still prevent the increase in the number of newly developed muscularized vessels. The conventional assessment showed in contrast a significant reduction of wall thickness and enlargement of the lumen in small and medium vessels in both models. These discrepancies between visual and digital analyses may reside in selecting representative fields of view by visual assessment, and/or may reflect the absence of consideration for non-muscularized vessels by the automatic digital analysis tool.

Regarding wall thickness, which is the key histological feature for PAH, automated digital quantification correlated well with that obtained with the conventional visual assessment (r = 0.80) as did the assessment of the lumen (r = 0.88). The two strong correlations lead to several considerations: 1-the current focus on muscular and semi-muscular vessels by the automated analysis introduces a very limited error in wall thickness measurements: as the same trends as the pathologist were found, intima and media measurements are expected to be accurate and 2-a partial assessment of rat lung section from randomly selected fields of view, such as provided by conventional visual assessment, allows an adequate quantification of the extent of vascular remodeling in PH animals.

Lastly, we verified whether our data correlated with the hemodynamic measurements and pulmonary functions, more specifically with mPAP and PVR. The correlations ranged from 0.71 to 0.83, in both models, suggesting that the measured wall thickening contributes to the elevation of mPAP and PVR.

Overall, these results validate the usefulness of a-SMA IHC staining to investigate the extent of PH severity. It also supports that the assessment of the muscularized vessels only, regardless of their nature, can be a surrogate marker of vascular remodeling during PH development. It also confirms the sensitivity and specificity of the fully validated software for the automated quantification of vascular remodeling in MCT-and SuHx-induced PH rats.

There are some limitations to this work, however: these data come from one experiment in each model, albeit involving several dozens of animals per rat model. Considering the inter-study variability in PH induction (29, 30), these results should be confirmed in a series of experiments for each model. In addition, the impact of Sildenafil on vascular remodeling is still debated; new clinical drugs with demonstrated anti-remodeling efficacy will provide a wider range of mechanisms to investigate. It would also be interesting to assess PH during its onset, and observe how models behave over their natural course. Importantly, this work focuses on the morphometric analysis of lung vessels but does not allow a complete histological assessment of PH, which should include an overall assessment of lung tissue, including inflammation, fibrosis, and oedema.

In summary, we have developed the first fully automated software for the assessment of vascular remodeling in the context of PH, enabling a faster and more comprehensive characterization of lung vessels. This tool provides an important aid to the pathologist in the global assessment of PH lung histology.

## Acknowledgments

We thank Yvon Julé for his support in manuscript editing and reviewing and Karine Bertotti and Damien Barbes for their initial work on the developed software.

## Author contributions

Conceptualization: CS, BL, CEL, MB, DS performed study concept and design. CS, MB, FC, and CEL performed animal studies and obtained in vivo data. BL and CS developed MorphoQuant, providing technical and material support. CS, BL, MB and CEL performed development of methodology. CS, FC and CEL performed the statistical analysis and data interpretation. CS, FC, CEL, and DS wrote and edited the manuscript. All authors read and approved the final manuscript.

## Data Availability Statement

The datasets used and/or analyzed during the current study are available from the corresponding author on reasonable request.

---

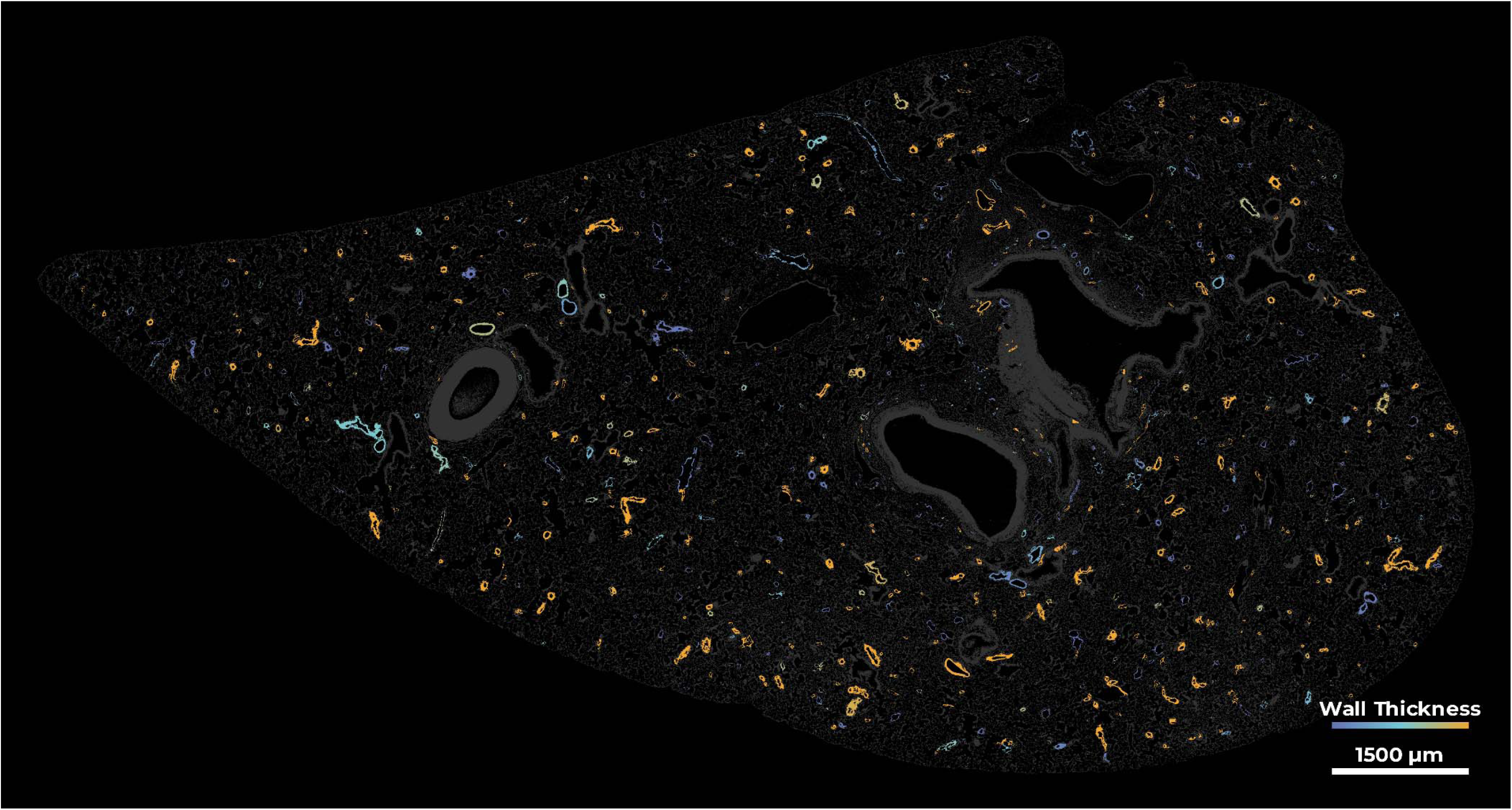
Suggestion for cover figure.

