## Supplemental figures for "UNRAVELING THE LUNG VASCULAR REMODELING IN PULMONARY HYPERTENSION USING A QUANTITATIVE DIGITAL PATHOLOGY SOFTWARE"

### SUPPLEMENTARY FIGURES

Fig. S1. Additional assessment of cardiac function and structure by echocardiography and invasive hemodynamics in MCT-induced PAH rats.

Fig. S2. Visual assessment of vessel proportion of muscular, semi-muscular and non-muscular vessels in MCT -induced PAH rats.

Fig. S3. Additional assessment of cardiac function and structure by echocardiography and invasive hemodynamics in SuHx-induced PAH rats.

Fig. S4. Visual assessment of vessel proportion of muscular, semi-muscular and non-muscular vessels in SuHx -induced PAH rats.

Fig. S5. Comparison and correlation of luminal radius between visual assessment and automated digital analysis in MCT- and SuHx-induced PAH rats.

Fig. S6. Comparison and correlation of vessel occlusion between visual assessment and automated digital analysis in MCT- and SuHx-induced PAH rats.

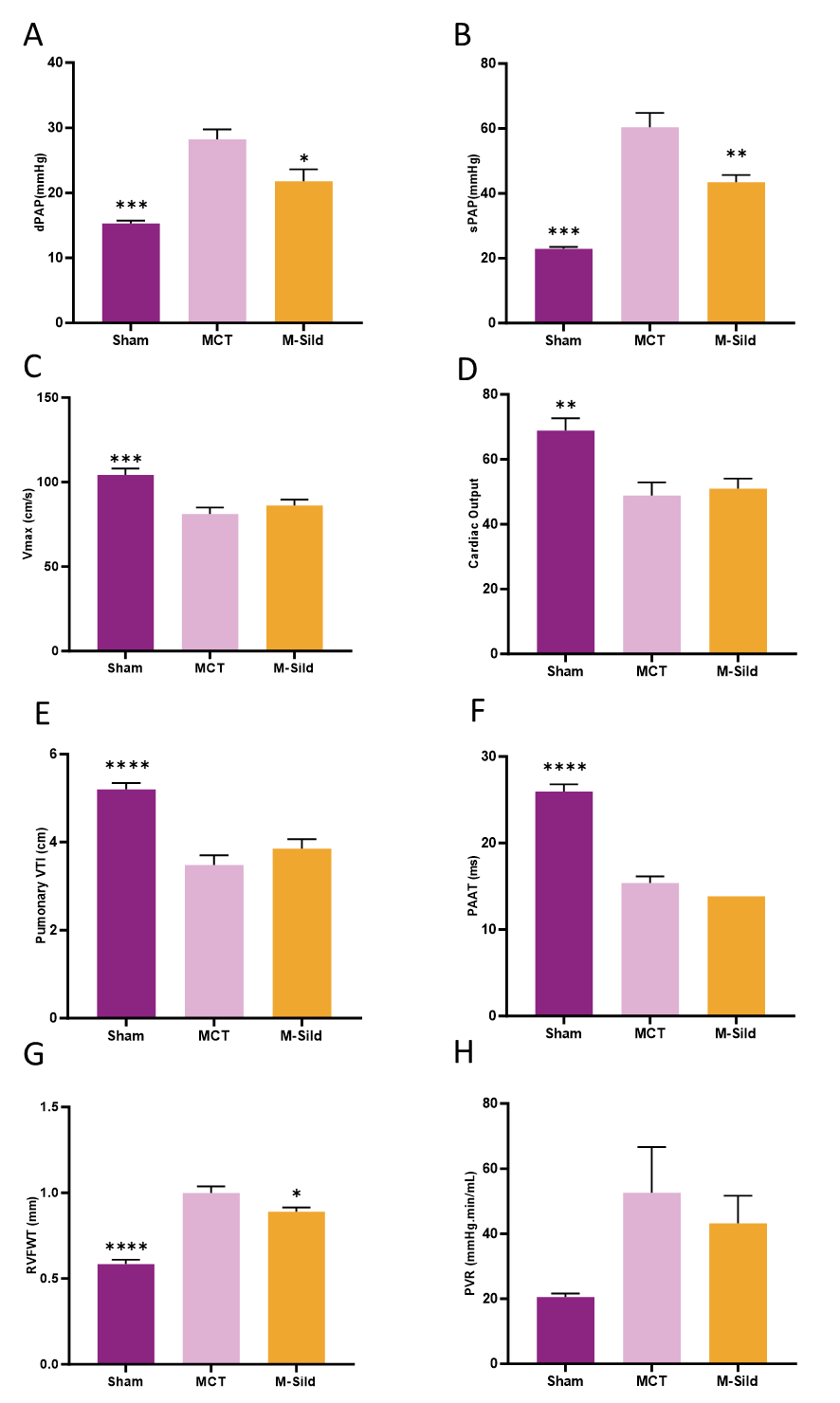

Fig. S1. Additional assessment of cardiac function and structure by echocardiography and invasive hemodynamics in MCT-induced PAH rats. (A) Diastolic pulmonary arterial pressure. (B) Systolic pulmonary arterial pressure. (C) Stroke Volume. (D) Cardiac output. (E) Pulmonary velocity time integral. (F) Pulmonary artery acceleration time. (G) Right ventricular free wall Thickness. (H) Pulmonary vascular resistance. The statistical analysis was performed with either a t-test or a Mann-Whitney to compare Sham or M-Sild to MCT. MCT: monocrotaline; Sild: Sildenafil. *: p < 0.05; **: p < 0.01; ***: p < 0.001; ****: p < 0.0001

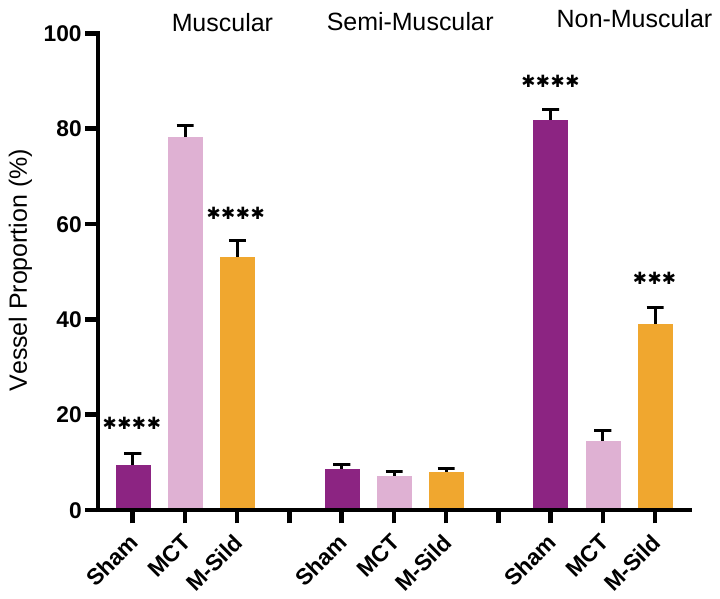

Fig. S2. Visual assessment of vessel proportion of muscular, semi-muscular and non-muscular vessels in MCT -induced PAH rats. The statistical analysis was performed with either a t-test or a Mann-Whitney to compare Sham or M-Sild to MCT. MCT: Monocrotaline; Sild: Sildenafil. *: p < 0.05; **: p < 0.01; ***: p < 0.001; ****: p < 0.0001.

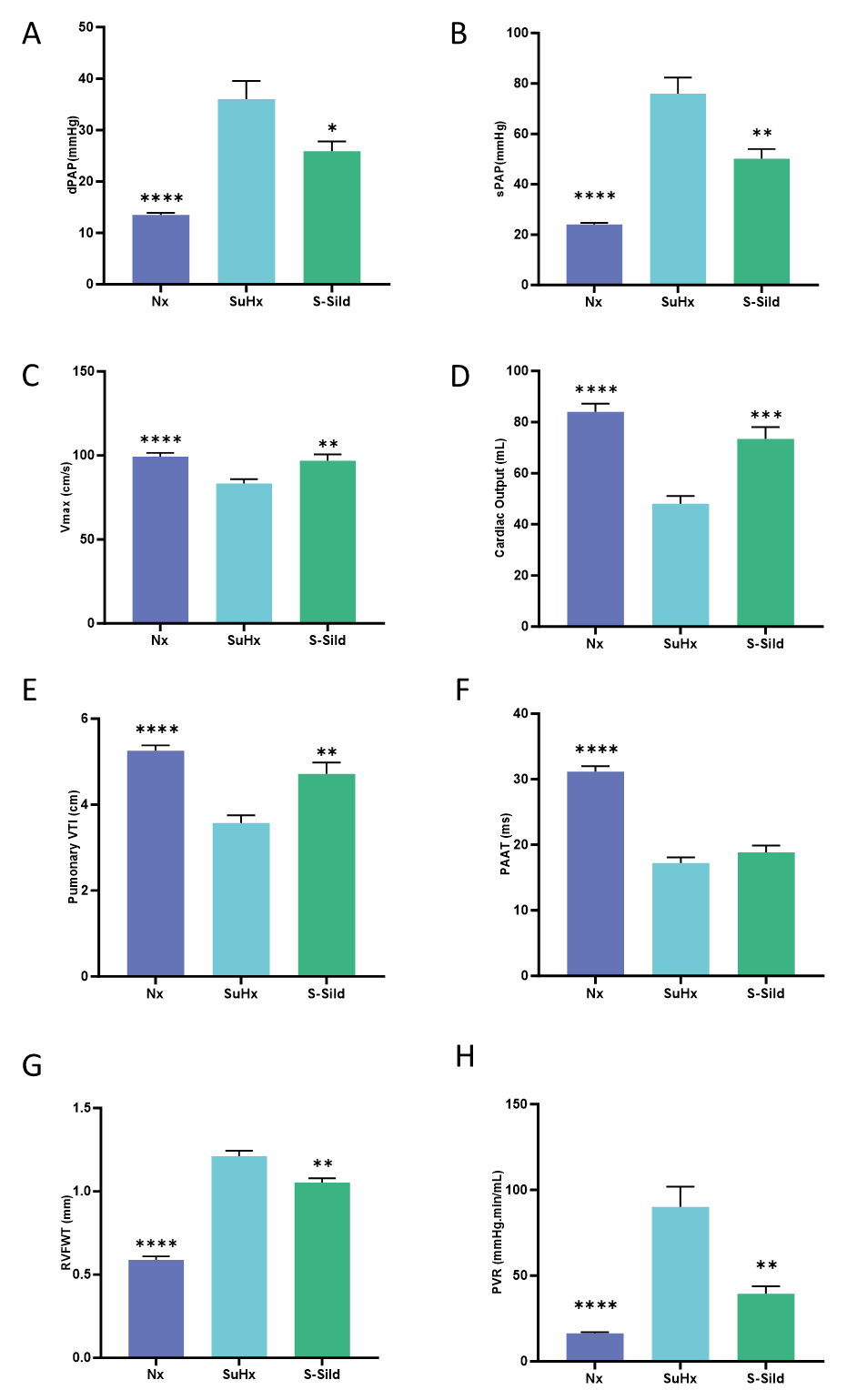

Fig. S3. Additional assessment of cardiac function and structure by echocardiography and invasive hemodynamics in SuHx-induced PAH rats. (A) Diastolic pulmonary arterial pressure. (B) Systolic pulmonary arterial pressure. (C) Stroke Volume. (D) Cardiac output. (E) Pulmonary velocity time integral. (F) Pulmonary artery acceleration time. (G) Right ventricular free wall thickness. (H) Pulmonary vascular resistance. The statistical analysis was performed with either a t-test or a Mann-Whitney to Nx or S-Sild to SuHx. Nx: Normoxic; SuHx: Sugen + hypoxia; Sild: Sildenafil. *: p < 0.05; **: p < 0.01; ***: p < 0.001; ****: p < 0.0001

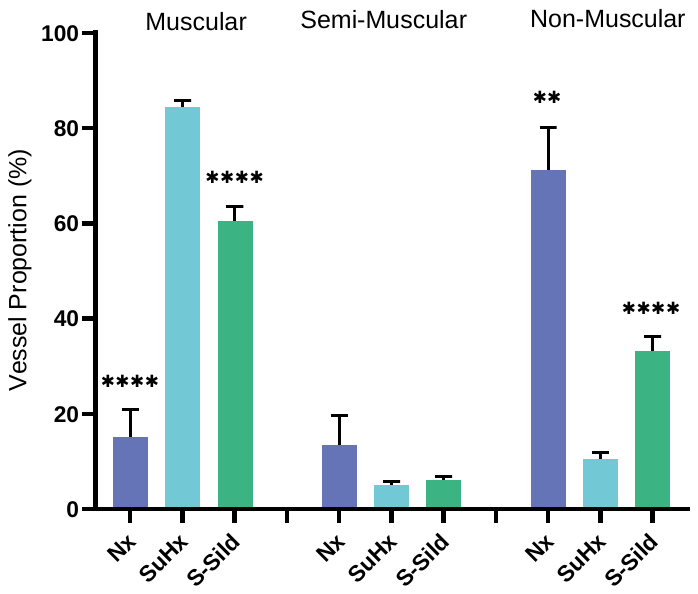

Fig. S4. Visual assessment of vessel proportion of muscular, semi-muscular and non-muscular vessels in SuHx -induced PAH rats. The statistical analysis was performed with either a t-test or a Mann-Whitney to compare SuHx or S-Sild to SuHx. SuHx: Sugen-hypoxia; Sild: Sildenafil.

*: p < 0.05; **: p < 0.01; ***: p < 0.001; ****: p < 0.0001.

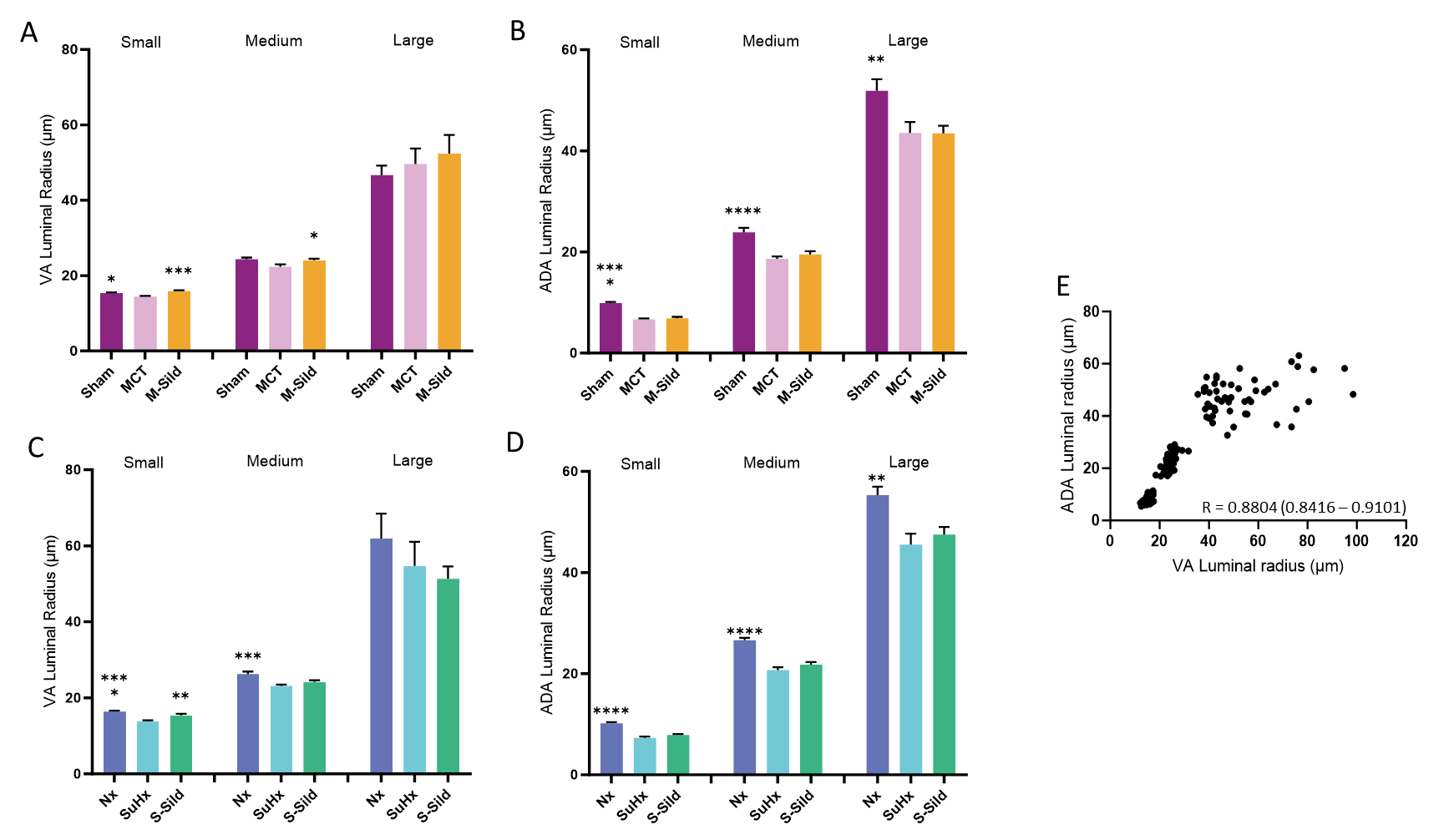

Fig. S5. Comparison and correlation of luminal radius between visual assessment and automated digital analysis in MCT- and SuHx-induced PAH rats. Luminal radius measured by visual assessment (VA) (A) and automated digital analysis (ADA) (B) in the MCT experiment. Luminal radius assessed by VA (C) and ADA (D) in the SuHx experiment. VA: visual assessment; ADA: automated digital analysis; Nx: Normoxic; SuHx: Sugen-hypoxia; MCT: monocrotaline; Sild: Sildenafil. *: p < 0.05; **: p < 0.01; ***: p < 0.001; ****: p < 0.0001. Comparison with either Vehicle groups

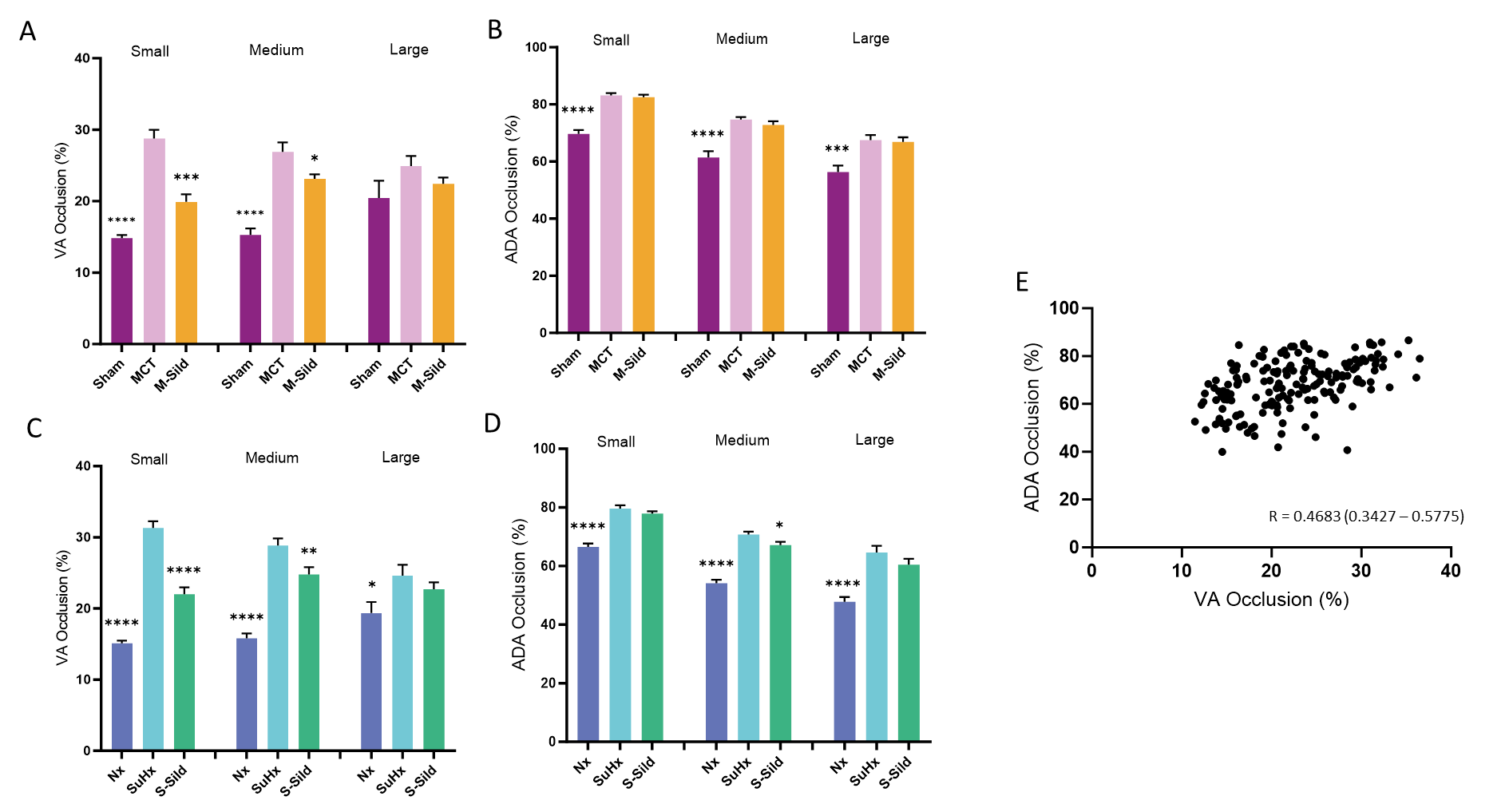

**Fig. S6**. Comparison and correlation of vessel occlusion between visual assessment and automated digital analysis in MCT- and SuHx-induced PAH rats. Vessel occlusion measured by visual assessment (VA) (A) and automated digital analysis (ADA) (B) in the MCT experiment. Vessel occlusion assessed by VA (C) and ADA (D) in the SuHx experiment. VA: visual assessment; ADA: automated digital analysis; Nx: Normoxic; SuHx: Sugen-hypoxia; MCT: monocrotaline; Sild: Sildenafil. *: p < 0.05; **: p < 0.01; ***: p < 0.001; ****: p < 0.0001. Comparison with either Vehicle groups.
